## Supplemental figures for "Sterol 14-α-demethylase is vital for mitochondrial functions and stress tolerance in *Leishmania major*"

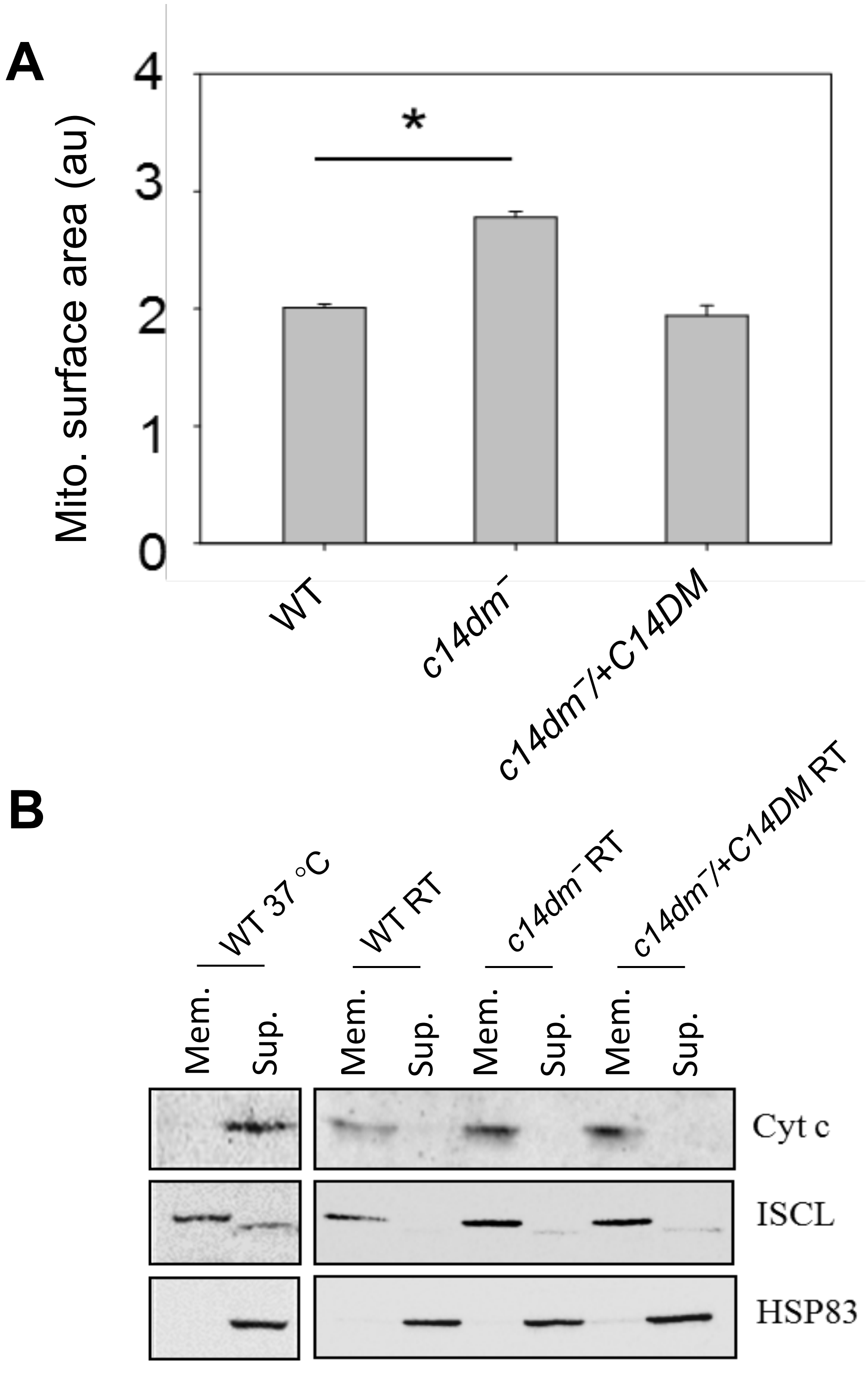


**Figure S1. *C14dm⁻* promastigotes show increased mitochondrial surface area but maintain mitochondrial membrane integrity at ambient temperature.** (**A**) After Mitotracker staining (Fig. 1A), the average mitochondrial surface areas in log phase promastigotes of WT, *c14dm⁻*, and *c14dm⁻/+C14DM* were determined using Image J (~200 cells were analyzed for each parasite line, au: arbitrary unit). (**B**) Log phase promastigotes were lysed with 0.035% of digitonin at 37 °C or room temperature (RT) and mitochondria enriched membrane fractions (Mem.) were separated from cytosolic fractions (Sup.) as described in Materials and Methods. Western blots were performed using antibodies against cytochrome *c*, ISCL (a mitochondrial membrane protein), and HSP83 (a cytosolic protein).

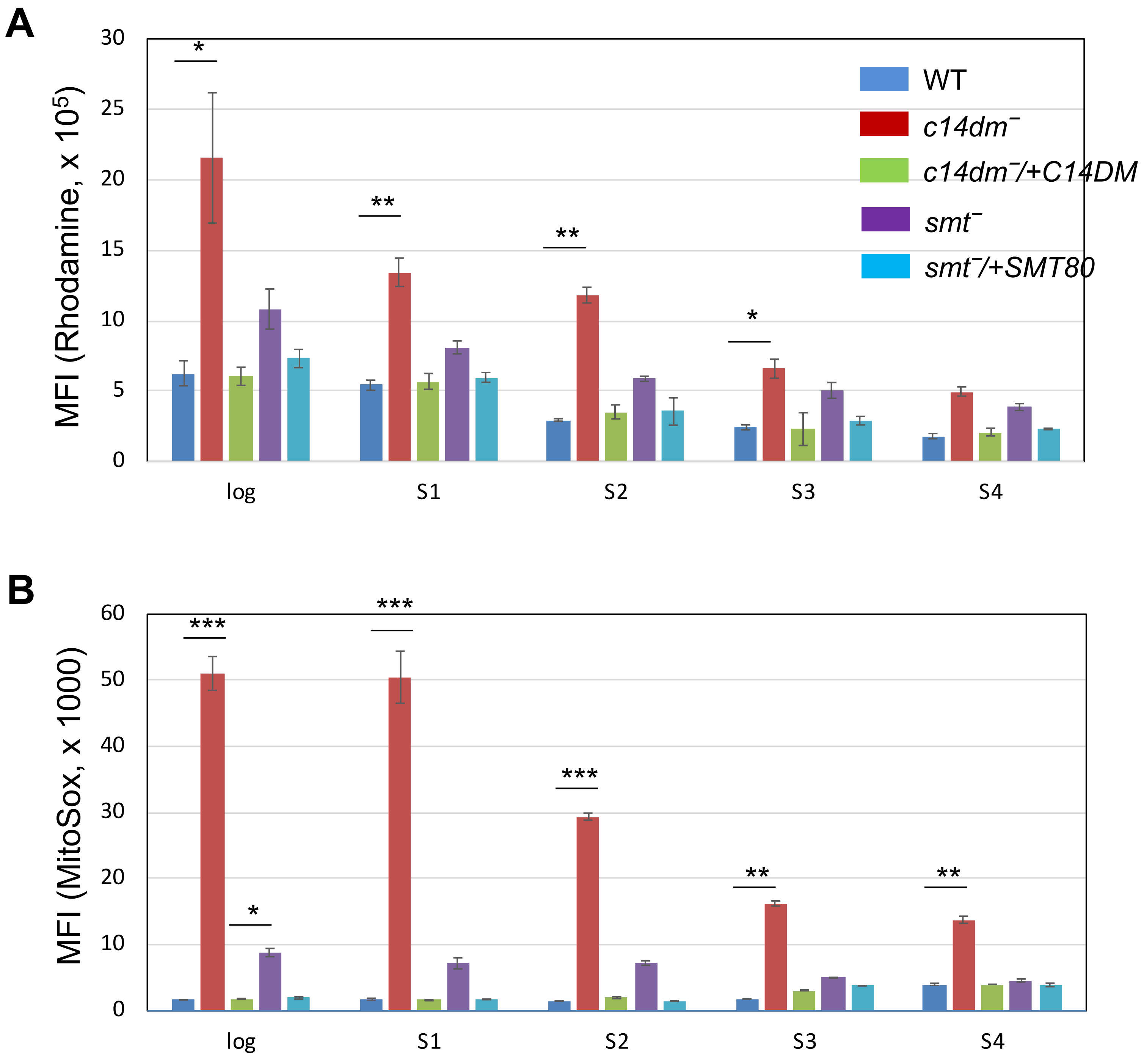


**Figure S2. *C14dm⁻* and *smt⁻* mutants show increased ΔMΨ and mitochondrial ROS.** Log phase and stationary phase (day 1-day 4) promastigotes were resuspended in PBS and labeled with 5 μg/ml of rhodamine 123 for 15 min (**A**) or with 5 μM of MitoSox Red for 25 min (**B**) at room temperature. MFIs were determined by flow cytometry. Error bars represent standard deviations from three independent experiments (***: *p*<0.001, **: *p*<0.01, *: *p*<0.05).

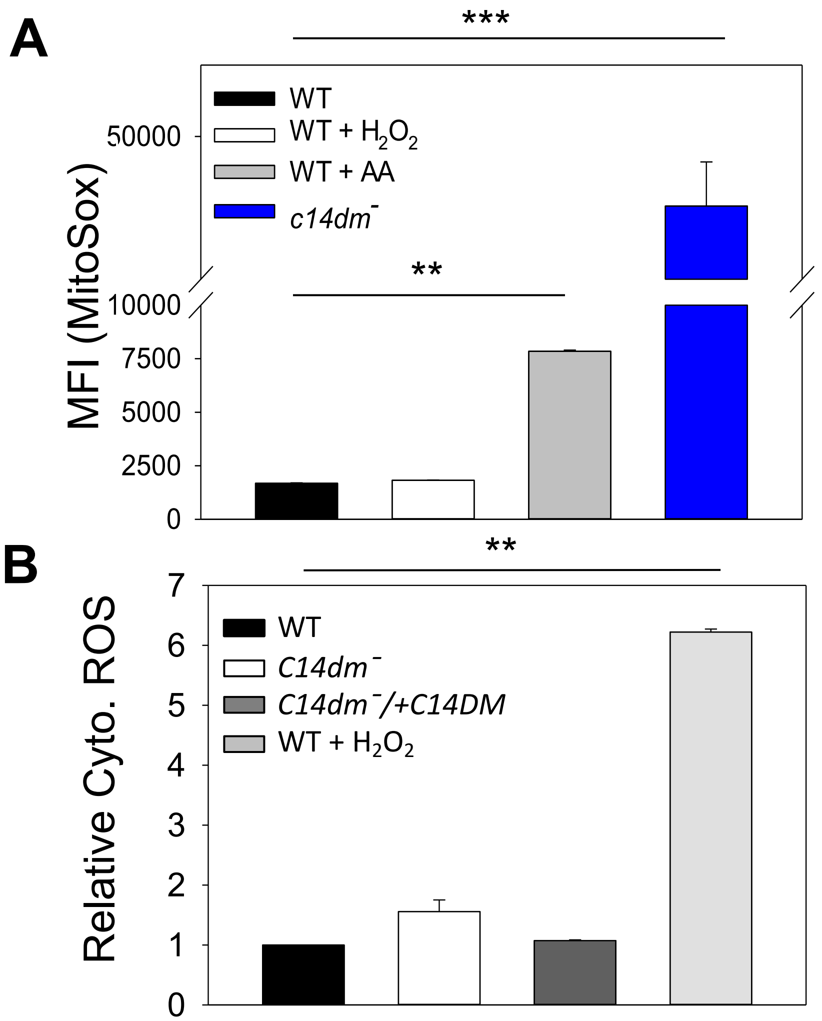


**Figure S3. *C14dm⁻* mutants accumulate ROS in the mitochondria.** Log phase promastigotes were labeled with 5 μM of MitoSox Red for 25 min (**A**) or 5 μM of DHE for 30 min (**B**) and MFIs were determined by flow cytometry. Effects of antimycin A (5 μM) and $H_2O_2$ (100 μM) on WT parasites were also monitored. Error bars represent standard deviations from three experiments.

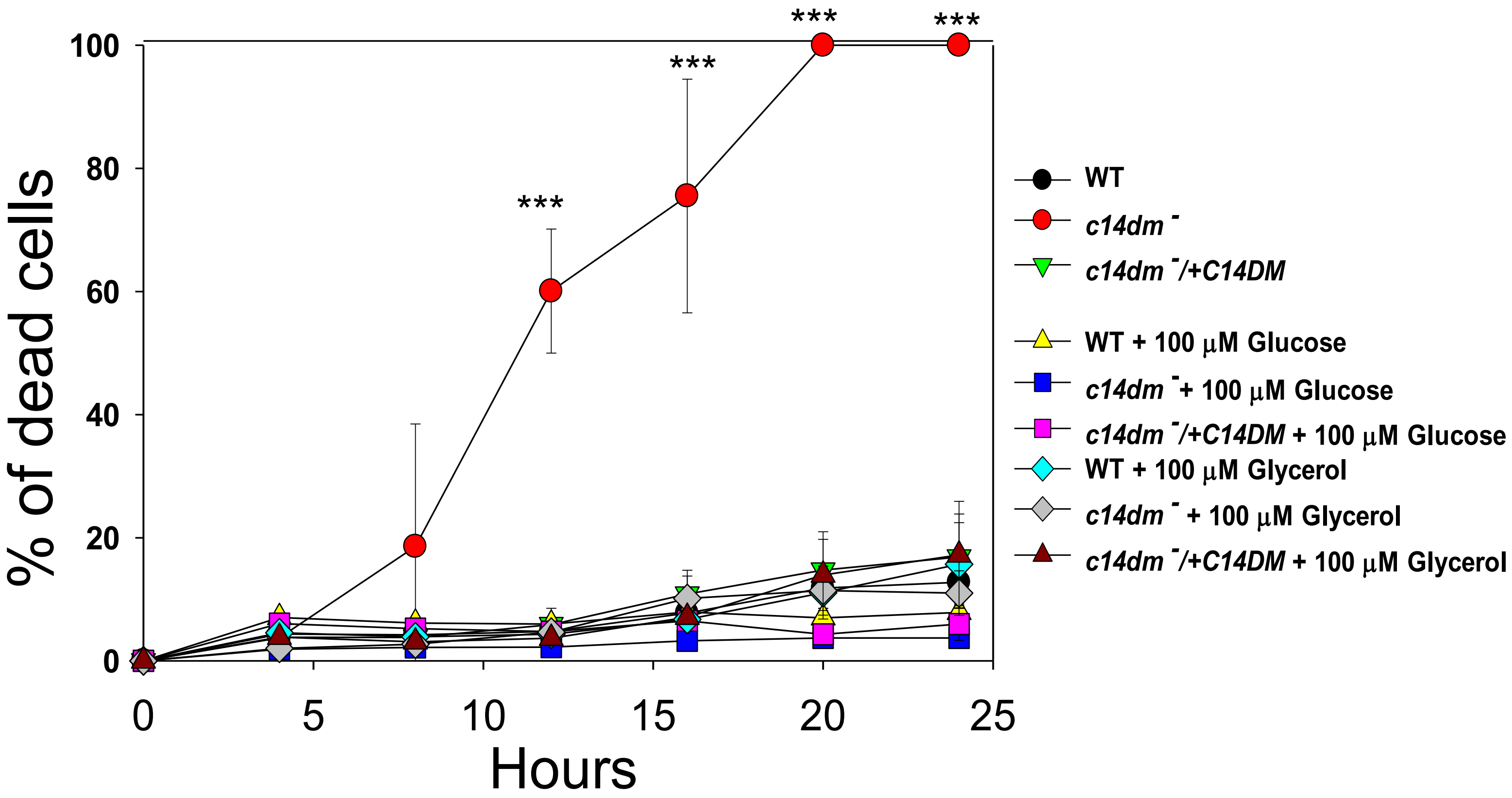


**Figure S4. 100 µM of glucose or glycerol is sufficient to rescue *c14dm*⁻ mutants in PBS.** Log phase promastigotes were incubated in PBS in the absence or presence of glucose (100 µM) or glycerol (100 µM) and percentages of dead cells were determined by flow cytometry at the indicated time points. Error bars represent standard deviations from three experiments.

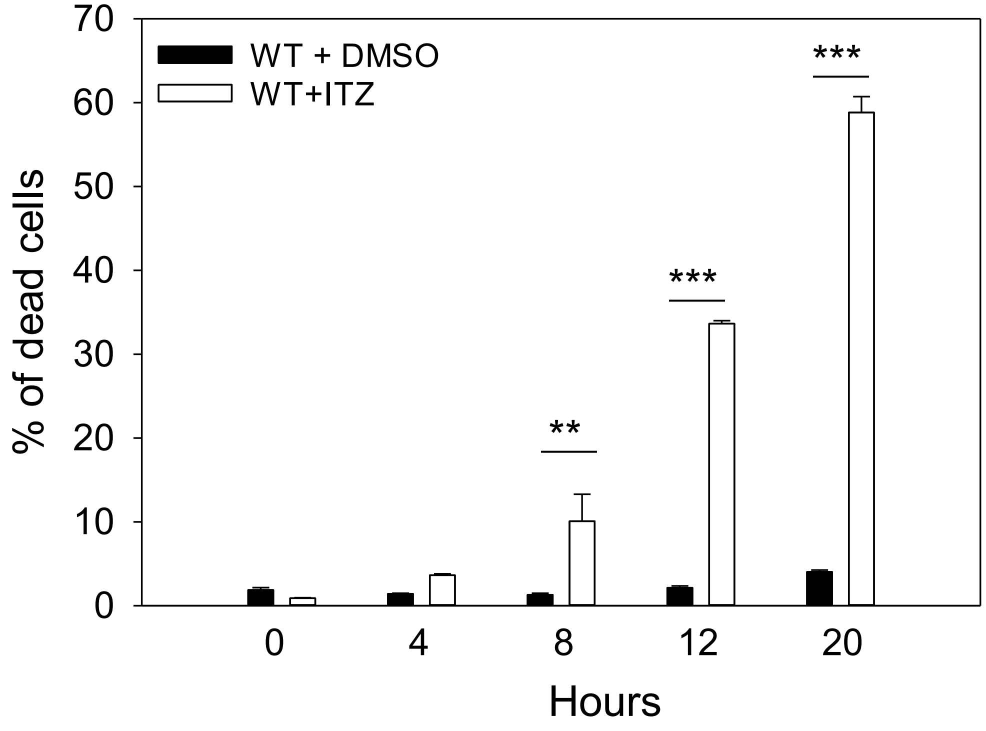


**Figure S5. ITZ-treated WT parasites are hypersensitive to glucose depletion.** (**A**) *L. major* WT promastigotes were cultivated in solvent alone (0.1% w/v of DMSO) or 0.2 μM of ITZ for 48 hours. Cells were then incubated in HBSS and percentages of dead cells were determined by flow cytometry at the indicated time points. Error bars represent standard deviations from three experiments (**: $p < 0.01$, ***: $p < 0.001$).

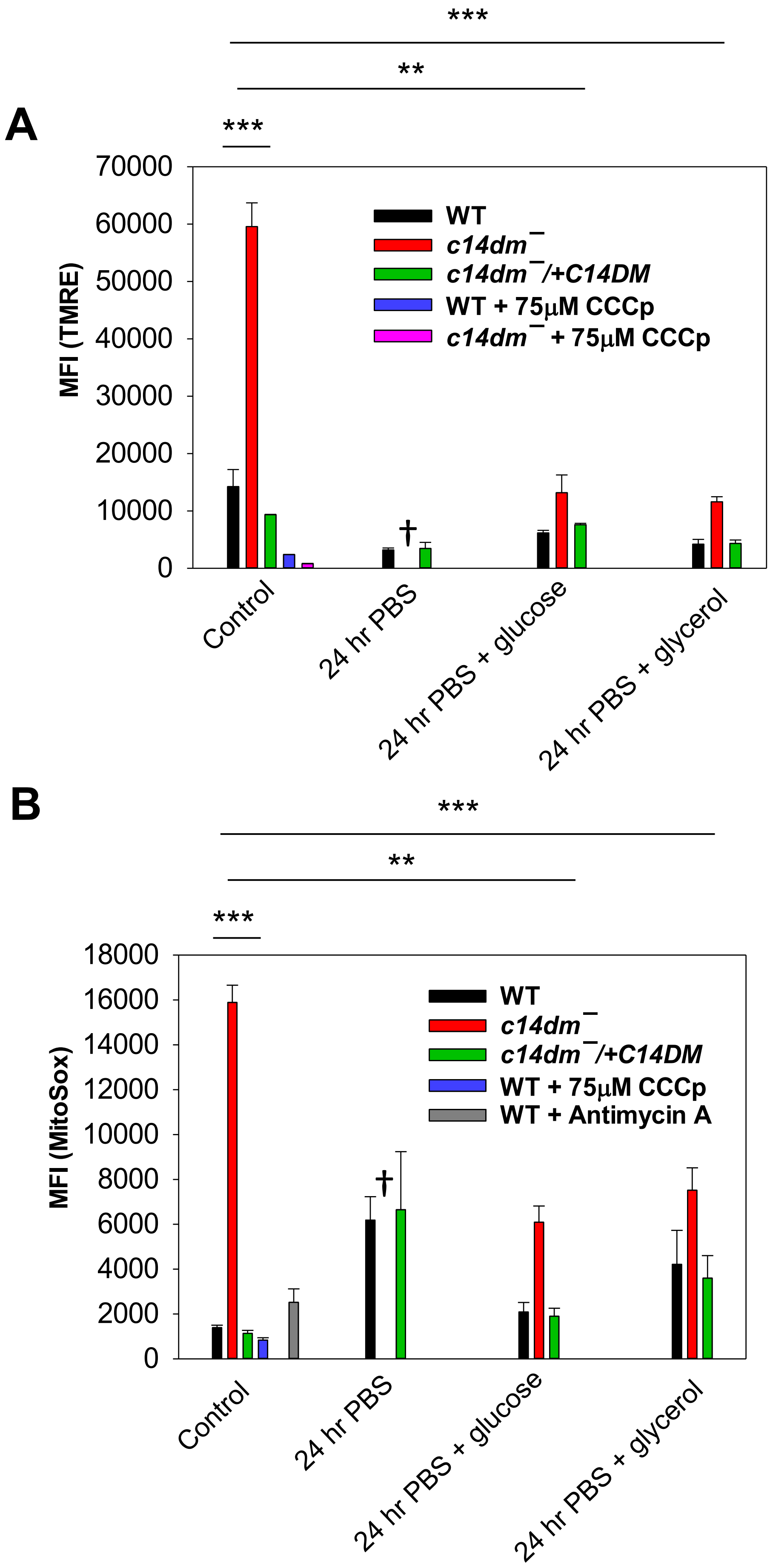


**Figure S6. *C14dm⁻* mutants rescued by glucose or glycerol show less mitochondrial defects.** Log phase promastigotes were incubated in PBS in the absence or presence of glucose (100 μM) or glycerol (100 μM) for 24 hours. ΔΦm (**A**) and mitochondrial ROS level (**B**) were determined by flow cytometry. Control cells represent cells analyzed at the beginning of incubation. †: no viable *c14dm⁻* mutants were available after 24 hours. Error bars represent standard deviations from three experiments (**: $p < 0.01$, ***: $p < 0.001$).
